# A Bias Correction Method for Substructure Estimation with Small Reference Sample Sizes

**DOI:** 10.64898/2026.11.04.763213

**Authors:** Souha Tifour, Adelle Price, Katie Marker, Audrey E. Hendricks

## Abstract

Reference data are important for accurate estimation of population substructure, yet sufficiently large reference samples for some populations are often unavailable. This limitation is especially important for methods that estimate population substructure from genetic summary data using reference allele frequencies derived from reference samples, such as Summix2. Here, we systematically evaluated how reference sample size for five continental reference groups [African (AFR), East Asian (EAS), European (EUR), Indigenous American (IAM), and South Asian (SAS)] affects Summix2 substructure estimates in simulated Latino-like, African American-like, and Peruvian-like populations, as well as real African American genetic data. We show that small reference sample sizes introduce systematic bias in substructure proportion estimates. We identified a sample size threshold of 150 individuals per reference group for accurate estimates with a mean estimation error below 0.05. We also developed two bias corrections: a simple inverse sample size correction and a flexible power-law correction. The power-law correction provided the greatest reduction in mean estimation error when all reference groups had small reference sample sizes, whereas the simple inverse sample size correction performed better when a single reference group had a small reference sample size. These findings provide practical guidance and bias adjustments for reliable population substructure estimation from genetic summary data, particularly when reference populations have small sample sizes.

## Introduction

Genetic summary data, including allele frequencies (AFs) and genome-wide association study (GWAS) summary statistics, are widely used because they enable secondary analyses without requiring access to individual-level genetic data including meta-analysis,^1^ Mendelian randomization,^2^ polygenic score development,^3^ and the incorporation of external control cohorts.^4,5^ However, genetic summary data aggregate information across individuals, masking individual-level heterogeneity such as population substructure. Heterogeneity is particularly important to model for genetically admixed populations, such as African American and Latino populations, but is also important to consider for within continental populations as well.^6^ Failure to account for this heterogeneity can bias downstream genetic analyses and limit their generalizability.^7^

Methods to estimate substructure from summary data exist. Summix^8^ and its extension, Summix2^7^ estimate substructure from AFs, Privé et al. developed a similar method using principal components from AFs,^9^ and ZMIX estimates substructure using AFs or Z scores.^10^ By estimating population substructure without requiring individual-level genotype data, these methods expand the utility of genetic summary data. However, because these methods rely on the reference group AFs to infer population substructure, their performance depends on the sample size and representativeness of the underlying reference groups.^7^

In practice, populations are represented unevenly across public genomic resources, resulting in substantial differences in reference sample sizes. Indigenous American (IAM) populations provide a clear example of this challenge. The 1000 Genomes Project (1KG)^11^ includes 347 individuals from admixed American (AMR) populations, compared with 661 African (AFR), 504 East Asian (EAS), 503 European (EUR), and 489 South Asian (SAS) individuals, but does not include a dedicated IAM reference group. Moreover, expanding IAM reference datasets must be considered within principles of Indigenous data sovereignty and community governance, meaning that larger reference samples may not always be available or appropriate.^12^

Here, we systematically evaluate how reference sample size affects population substructure estimation in summary data using Summix2. We identify a simple threshold to ensure minimal estimation error (<5%) and further develop methods to correct for bias from small reference sample sizes, which we assess in a variety of simulation scenarios and real genetic summary data.

## Materials and Methods

### Population Substructure Estimation

Summix^8^ estimates mixing proportions (*π*_*k*_) in genetic summary data by minimizing the least-squares difference between the observed AFs (*AF*_obs,*j*_) and a weighted combination of AFs from K reference groups (*AF*_ref,*j,k*_) across *M* single nucleotide polymorphisms (SNPs), where *j* indexes SNPs and *k* indexes the reference groups (Equation 1). Optimization is performed using sequential quadratic programming (SQP), subject to the constraints that mixing proportions are non-negative and sum to one. Summix2,^7^ available through Bioconductor, builds on Summix by enabling local and fine scale population substructure estimation and includes additional methodological improvements. In this study, we used Summix2 to estimate mixing proportions from simulated and real genetic summary data.

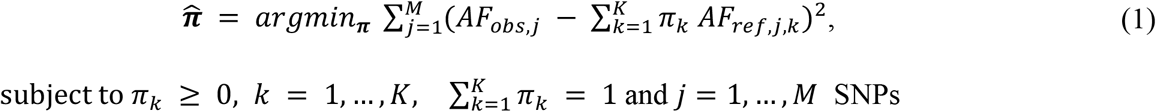

To reduce bias in Summix2 mixing proportion estimates arising from small reference sample sizes, we developed two bias correction methods: a simple 1/*N* correction and a flexible power-law *a*/*N*^*b*^ correction.

Let 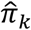 denote the mixing proportion estimated by Summix2 for reference group *k, π*_*k*_ the true mixing proportion for reference group *k*, and *N*_*k*_ the reference sample size for reference group *k*. Bias was defined as

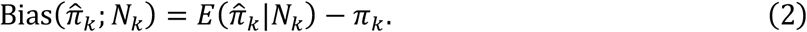

### Simple bias correction

For the simple correction, we modeled the magnitude of bias as inversely proportional to reference sample size:

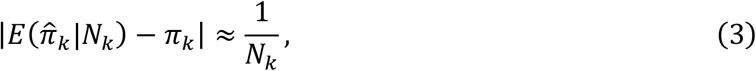

Allowing for positive or negative bias, Equation (3) can be expressed as,

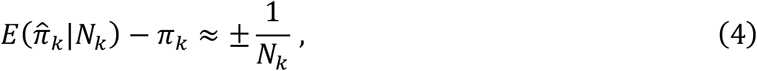

Rearranging Equation (4) to express the true mixing proportion, *π*_*k*_, gives:

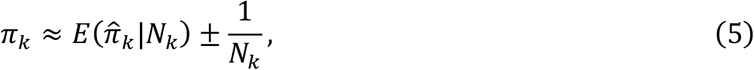

Based on Equation (5), we defined the bias-corrected mixing-proportion estimate, 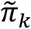, by applying the correction to 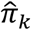. The direction of the adjustment was specified empirically using 1/*K* as a threshold, where *K* is the total number of reference groups. Specifically, 1/ *N*_*k*_ was subtracted when 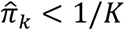, added when 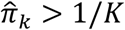, and no adjustment was made when 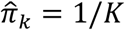:

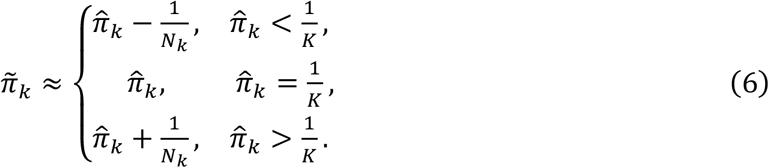

### Flexible bias correction

To allow the magnitude and rate of decay of bias to vary with reference sample size, we modeled the magnitude of bias using a power-law^13^ relationship:

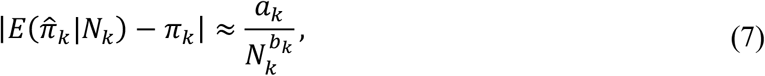

where *a*_*k*_ is a scale parameter and *b*_*k*_ is the exponent governing the rate at which bias decreases with increasing reference sample size. Accounting for positive and negative bias and solving for *π*_*k*_, Equation (7) gives

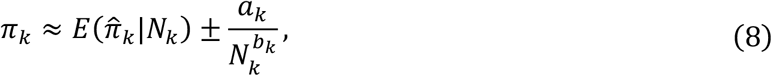

Using the same empirically determined direction of bias as for the simple 1/*N* correction, the power-law correction was applied as:

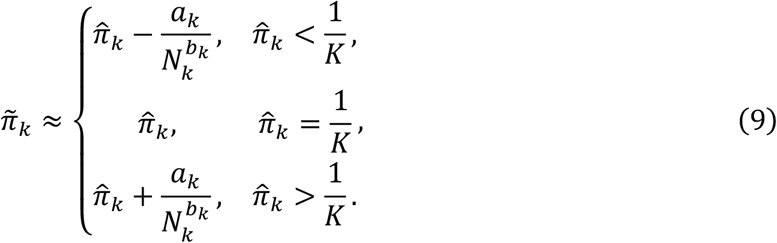

To estimate *a*_*k*_ and *b*_*k*_, we generated 100 sets of mixing proportions for parameter estimation by sampling across the five reference groups from a symmetric Dirichlet distribution with concentration parameters *α* = (1,1,1,1,1). Simulations were performed for reference sample sizes *N* = 1, …,1000, with 1,000 replicates per reference sample size, following the procedure described in the Simulation Study section. For each reference group, set of mixing proportions, and reference sample size, estimation error, 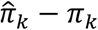, was calculated for each replicate and averaged across the 1,000 replicates to estimate bias. The absolute value of the estimated bias was fitted to the power-law function 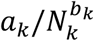 using nonlinear least squares separately for each reference group and set of mixing proportions. Fits that failed to converge were excluded, and the resulting estimates of 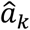 and 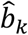 were summarized across successfully fitted sets of mixing proportions using the median for each reference group.

### Post-correction constraints

For both correction methods, negative corrected estimates were truncated to zero, and the complete vector of mixing proportions was renormalized to sum to one:

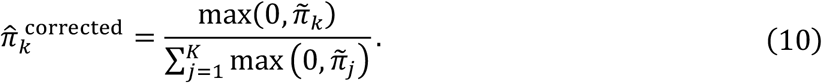

If truncation resulted in an all-zero vector, making the denominator in Equation (10) zero, the final corrected mixing proportions were set to 1/*K* for all reference groups.

When all reference groups were varied jointly, the correction was applied directly to all reference groups. When a single reference group was varied, the correction was applied directly only to the varying reference group followed by normalization across all reference groups.

### Data

This study used observed AFs from gnomAD v3.1.2 ^14^ and reference AFs from the 1KG and Human Genome Diversity Project (HGDP)^15^, all aligned to GRCh38. The gnomAD v3.1.2 dataset included 644,267,978 variants that passed gnomAD quality control (QC) from 76,156 whole genomes across ten population groups [African/African American (AFR/AFRAM), Amish (AMI), Latino/Admixed American (AMR), Ashkenazi Jewish (ASJ), East Asian (EAS), Finnish (FIN), non-Finnish European (NFE), Middle Eastern (MID), South Asian (SAS), and Other (OTH)] and served as the observed data. Reference AFs were obtained for 76 populations from HGDP and 1KG and harmonized with gnomAD v3.1.2. We merged the observed gnomAD data and 1KG and HGDP reference data by chromosome, variant position, reference allele, and alternate allele. Variants with missing values in either dataset were removed. Variants with a minor allele frequency (MAF) greater than 1% in at least one finer-scale reference population or one gnomAD population group were retained, resulting in 75,526,117 variants across the autosomes. Continental reference AFs for African (AFR), European (EUR), South Asian (SAS), East Asian (EAS), and Indigenous American (IAM) were derived from 50 finer-scale reference populations, as described further in Stoneman et al.^7^

### Simulation Study

Reference AFs were simulated independently for each of the five continental reference groups at sample sizes of *N* = 5, 10, 25, 35, 45, 50, 75, 100, 150, 250, 500, 750, and 1000. For each group and sample size, allele counts were sampled from 2N chromosomes according to the corresponding continental reference AFs using the `rmultinom` function in base R. Simulated reference AFs were then calculated by dividing the sampled allele counts by (2N).

Observed AFs were simulated for three admixed populations with distinct mixing proportions reported by Arriaga-MacKenzie et al^8^: Latino-like (AFR-like 4.4%, EUR-like 43.8%, IAM-like 46.9%, EAS-like 4.9%, SAS-like 0.0%), African American-like (AFR-like 85.2%, EUR-like 14.8%, IAM-like 0.0%, EAS-like 0.0%, SAS-like 0.0%), and Peruvian-like (AFR-like 2.7%, EUR-like 19.6%, IAM-like 76.8%, EAS-like 0.9%, SAS-like 0.0%). For each simulated population, SNP genotypes were generated within each reference group using the `rmultinom` function in base R under Hardy-Weinberg equilibrium, with genotype probabilities determined by the corresponding reference AFs. Genotype counts were then combined across reference groups according to the specified mixing proportions to generate AFs for the simulated observed population. The observed population sample size was fixed at 10,000 individuals across all reference sample sizes and admixed populations. For each setting, 1,000 simulation replicates were generated using the 99,187 SNPs from chromosome 22 that remained after the filtering described above. Additional details and code are provided in the accompanying GitHub repository (see Data and Code Availability). We evaluated three reference sample size scenarios:

- **<u>Scenario 1 (reference sample sizes varied together)</u>**: Reference sample sizes were varied jointly across all five reference groups, with each reference group assigned the same sample size. The sample sizes evaluated were *N* = {5, 10, 25, 35, 45, 50, 75, 100, 150, 250, 500, 750, 1000}.
- **<u>Scenario 2 (one reference group varied, others fixed at 500)</u>**: The reference sample size for one reference group was varied across *N* = {5, 10, 25, 35, 45, 50, 75, 100, 150, 250, 500, 750, 1000}, while the reference sample sizes for the remaining four reference groups were fixed at *N* = 500.
- **<u>Scenario 3 (one reference group varied, others fixed at 150)</u>**: The reference sample size for one reference group was varied across *N* = {5, 10, 25, 35, 45, 50, 75, 100, 150, 250, 500, 750, 1000}, while the reference sample sizes for the remaining four reference groups were fixed at *N* = 150.

In Scenarios 2 and 3, the IAM reference sample size was varied for the Latino-like and Peruvian-like populations, and the AFR reference sample size was varied for the African American-like population.

For each continental reference group and simulation scenario, estimation error was calculated as the difference between the estimated and true mixing proportions:

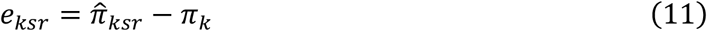

where *π*_*k*_ denotes the true mixing proportion for continental reference group *k*, and 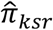. denotes the corresponding Summix2 estimate in replicate *r* under simulation scenario *s*. Bias was estimated as the mean estimation error across the *R* = 1,000 simulation replicates:

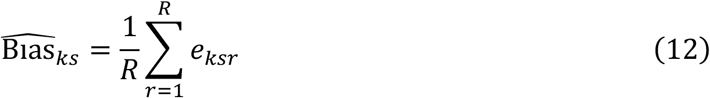

To quantify the magnitude of bias independent of direction, we calculated mean absolute estimation error (MAE) by averaging absolute estimation error across the *R* = 1,000 simulation replicates:

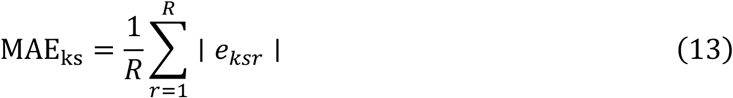

To quantify overall estimation error across the five continental reference groups, mean total absolute estimation error was calculated across the *R* = 1,000 simulation replicates as

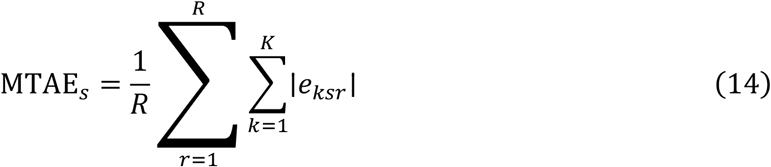

### Real-data evaluation

We evaluated the bias correction methods using real genetic summary data. Observed AFs were obtained from chromosome 22 gnomAD v3.1.2 AFR/AFRAM summary data, and reference AFs were obtained from harmonized 1KG/HGDP data for the AFR, EUR, EAS, SAS, and IAM reference groups. To evaluate the effect of reference sample size, individuals from each reference group were subsampled to generate reference AFs at different sample sizes, while the observed gnomAD AFs were held fixed. For each evaluated reference sample size, the subsampled reference AFs were merged with the observed gnomAD AFs by chromosome, variant position, reference allele, and alternate allele, and Summix2 was rerun using the resulting reference AFs.

Because the true mixing proportions are unknown in real data, estimates obtained from large reference sample sizes served as the benchmark. For the AFR, EUR, EAS, and SAS reference groups, the benchmark was defined using reference panels with *N* = 500. For the IAM reference group, which comprised 46 individuals, the benchmark was defined using *N* = 45, the largest rounded sample size.

We evaluated two real-data settings designed to parallel the simulation scenarios:

- **<u>Setting 1 (reference sample sizes varied together where available)</u>**: all five reference groups had the same reference sample size for *N* = 5,15,25,35, and 45. For larger evaluated reference sample sizes (*N* = 75,100,150,250, and 500), the IAM reference sample size remained fixed at *N* = 45, while the AFR, EUR, EAS, and SAS reference sample sizes increased.
- **<u>Setting 2 (one reference group varied, others fixed at benchmark)</u>**: one continental reference group varied over *N* ∈ {5, 15, 25, 35, 45, 75, 100, 150, 250, 500}, while the remaining reference groups were fixed at the benchmark sample sizes (*N* = 500 for AFR, EUR, EAS, and SAS reference groups and *N* = 45 for the IAM reference group).

For each setting, we compared uncorrected Summix2 estimates with bias-corrected estimates. Performance was evaluated using the absolute deviation of the mixing-proportion estimate from the large reference sample benchmark for each continental reference group, and the total absolute deviation summed across all reference groups.

## Results

### Total absolute estimation error decreases approximately inversely with reference sample size

In Scenario 1, in which all five reference groups had the same reference sample size, total absolute estimation error showed an approximately inverse relationship with reference sample size across the African American-like, Latino-like, and Peruvian-like simulated admixed populations (Figure 1). Mean total absolute estimation error decreased from 0.499–0.521 at *N* = 5 to 0.027–0.043 at *N* = 150, with mean total absolute estimation error below 0.05 for all three simulated admixed populations at *N* = 150. These results show that jointly limiting the reference sample sizes of all five reference groups increases total estimation error in Summix2 mixing-proportion estimates.

**Figure 1.**
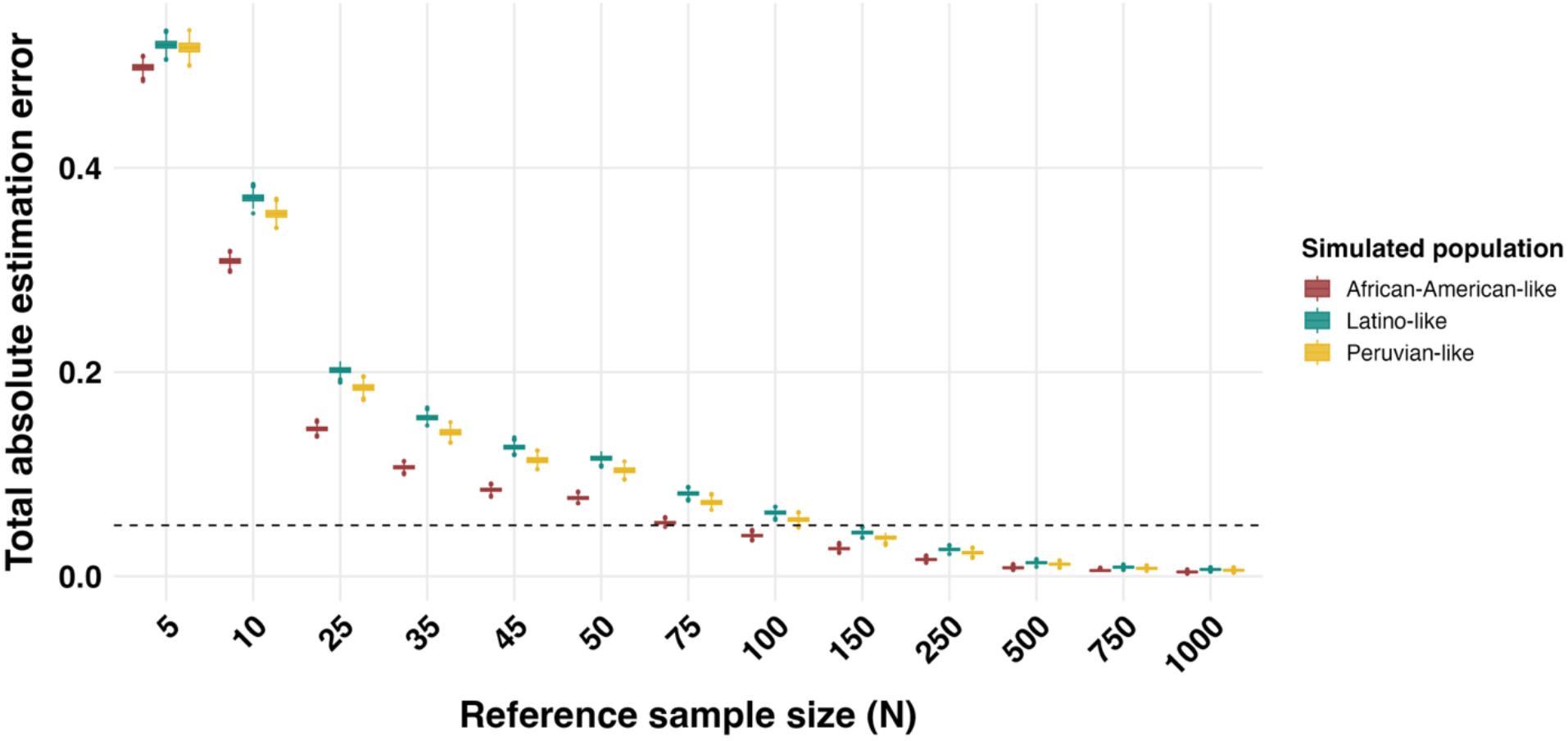
Effect of reference sample size on total absolute estimation error in Scenario 1. All five reference groups (AFR, EAS, EUR, IAM, and SAS) were simulated with the same reference sample size (*N*). Total absolute estimation error was calculated across the five reference groups for each simulation replicate for the African American-like, Latino-like, and Peruvian-like simulated admixed populations. Boxplots show the distribution of total absolute estimation error across 1,000 simulation replicates at each reference sample size. The dashed horizontal line indicates a total absolute estimation error of 0.05.

### Small sample size of one reference group increases estimation error in other reference groups

In Scenario 2, the reference sample size of one reference group was varied while those of the other four reference groups were fixed at *N* = 500. For the Latino-like simulated admixed population, IAM was the varying reference group, while AFR, EAS, EUR, and SAS were fixed reference groups (Figure 2). At an IAM reference sample size of *N* = 5, mean absolute estimation error was 0.179 for IAM and 0.143 for EAS, despite the EAS reference sample size remaining fixed at *N* = 500 (Figure 2A–B). As the IAM reference sample size increased from *N* = 5 to *N* = 150, mean absolute estimation error decreased to 0.009 for IAM and 0.007 for EAS. Mean total absolute estimation error across the five reference groups decreased from 0.380 to 0.024 over the same range of IAM reference sample sizes (Figure 2C). These results show that limiting the reference sample size of one reference group can increase estimation error in other reference groups even when their reference sample sizes remain fixed. Results for the African American-like and Peruvian-like simulated admixed populations and for Scenario 3, in which the four fixed reference groups had reference sample sizes of *N* = 150, are presented in the Supplementary Results (Figures S2–S3)

**Figure 2.**
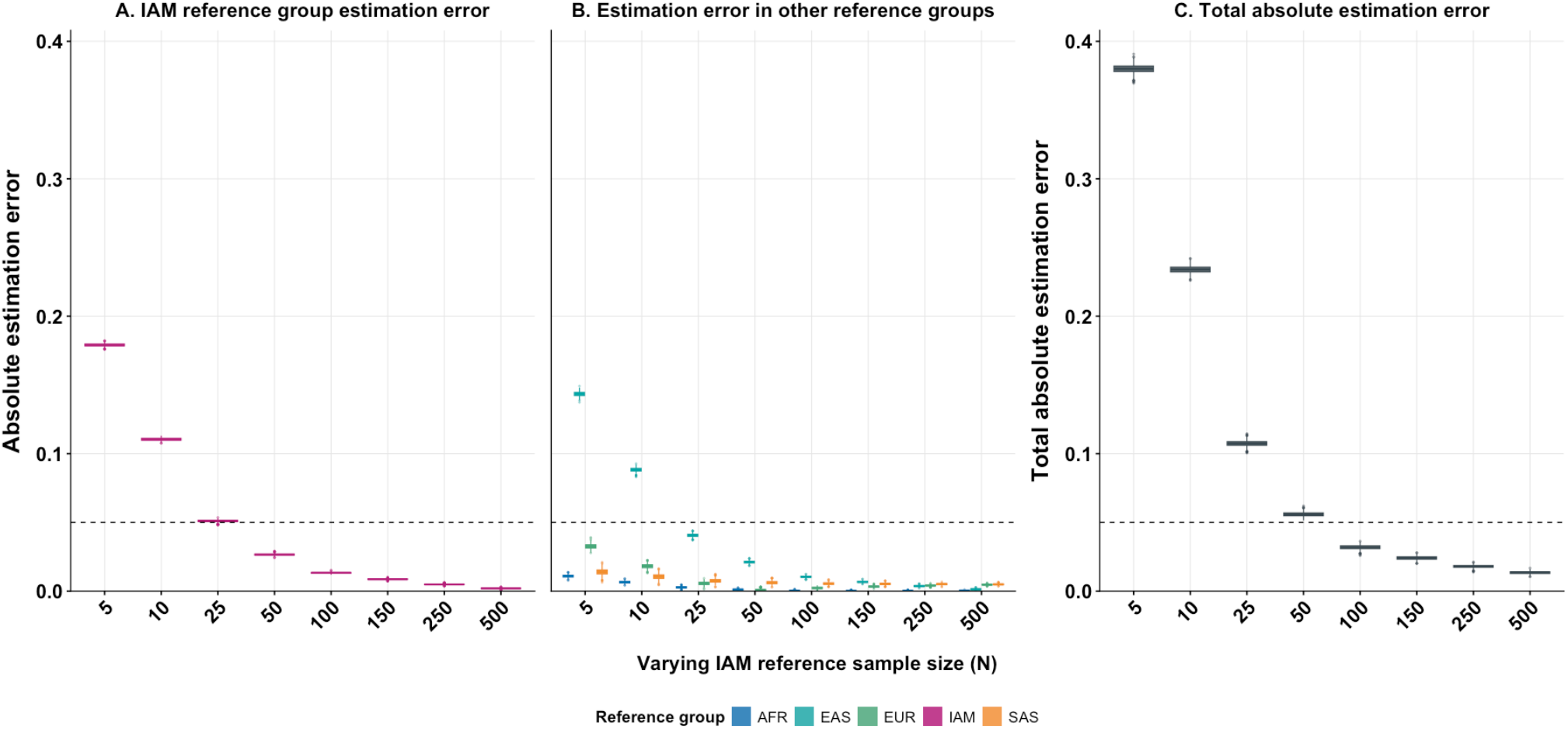
Effect of limiting the reference sample size of one reference group on absolute estimation error in Scenario 2. For the Latino-like simulated admixed population, IAM was the varying reference group, while AFR, EAS, EUR, and SAS were fixed at a reference sample size of *N* = 500. (A) Absolute estimation error for the varying reference group (IAM). (B) Absolute estimation error for the fixed reference groups (AFR, EAS, EUR, and SAS) as the IAM reference sample size increased. (C) Total absolute estimation error across all five reference groups. Boxplots show the distributions of absolute estimation error across 1,000 simulation replicates at each IAM reference sample size. The dashed horizontal line indicates 0.05.

### Bias correction reduces estimation error at small reference sample sizes

The empirical relationship between reference sample size and estimation error observed in the simulation analyses motivated the development of two corrections: the 1/*N* correction and the power-law *a*/*N*^*b*^ correction. We evaluated whether these corrections reduced total absolute estimation error in Summix2 mixing proportion estimates across the three simulation scenarios (Figure 3). Both corrections produced their largest reductions in mean total absolute estimation error at small reference sample sizes, while differences between corrected and uncorrected estimates decreased as reference sample size increased.

**Figure 3.**
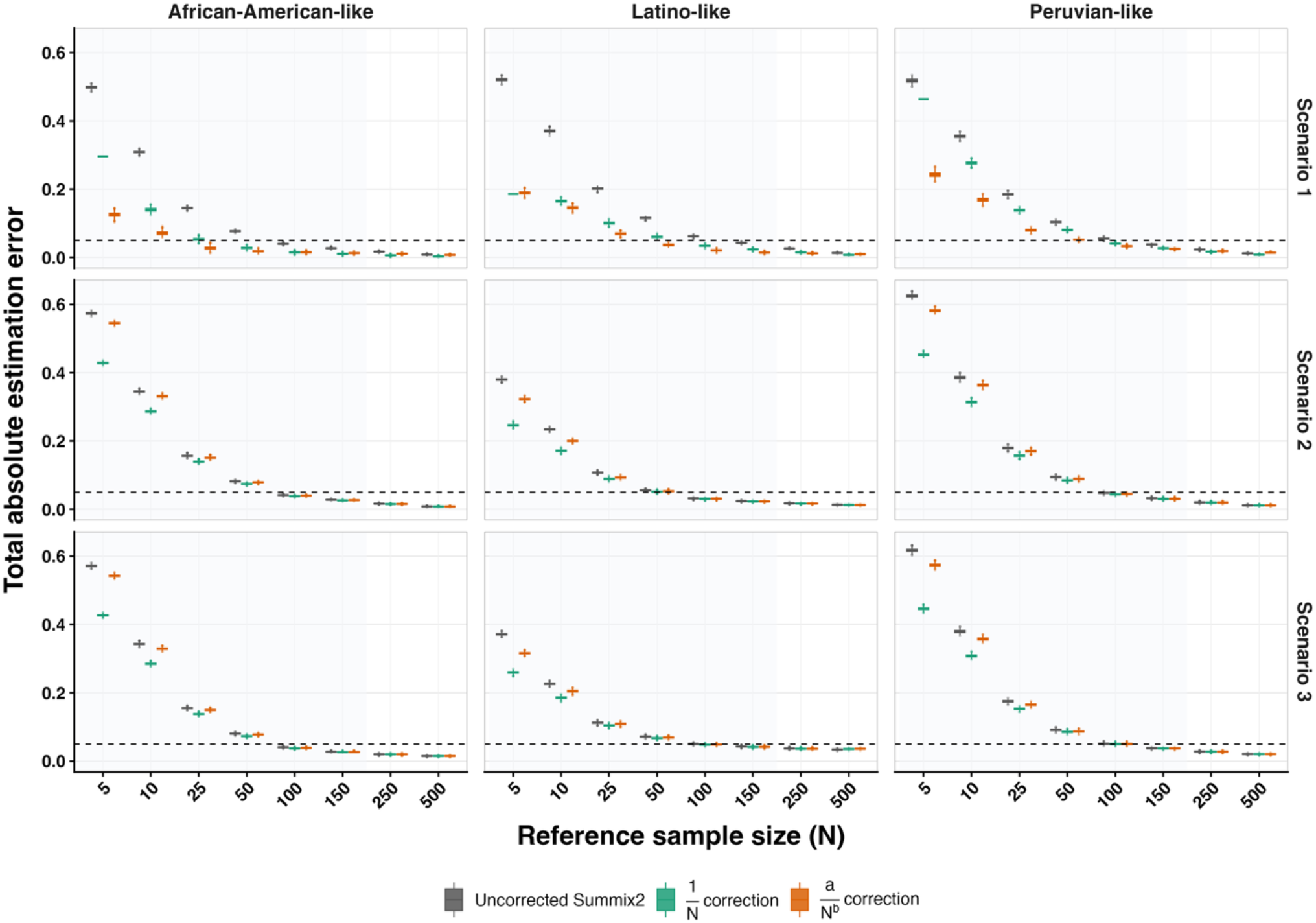
Performance of reference-sample-size corrections across simulation scenarios. Total absolute estimation error is shown for uncorrected Summix2 estimates and estimates after application of the 1/*N* and power-law *a*/*N*_*k*_ corrections for the African American-like, Latino-like, and Peruvian-like simulated admixed populations (columns). In Scenario 1, reference sample sizes were varied jointly across all five reference groups, with all five reference groups assigned the same reference sample size at each evaluated *N*. In Scenarios 2 and 3, the varying reference group was AFR for the African American-like population and IAM for the Latino-like and Peruvian-like populations; the reference sample sizes of the other four reference groups were fixed at *N* = 500 in Scenario 2 and *N* = 150 in Scenario 3 (rows). Boxplots show the distribution of total absolute estimation error across 1,000 simulation replicates at each reference sample size. The shaded region indicates reference sample sizes *N* ≤ 150, and the dashed horizontal line indicates a total absolute estimation error of 0.05.

In Scenario 1, in which reference sample sizes were varied jointly across all five reference groups, the power-law *a*/*N*^*b*^ correction generally produced lower mean total absolute estimation error than the 1/*N* correction at small reference sample sizes across the three simulated admixed populations (Figure 3, top row). For example, for the African American-like simulated admixed population at *N* = 5, mean total absolute estimation error decreased from 0.499 for the uncorrected estimates to 0.296 with the 1/*N* correction and 0.125 with the power-law correction. At *N* = 25, mean total absolute estimation error was 0.145 for the uncorrected estimates, 0.053 with the 1/*N* correction, and 0.028 with the power-law *a*/*N*^*b*^ correction.

In Scenarios 2 and 3, in which the reference sample size of one reference group varied while those of the other four reference groups were fixed at *N* = 500 and *N* = 150, respectively, the 1/*N* correction consistently produced the lowest mean total absolute estimation error across the three simulated admixed populations (Figure 3, middle and bottom rows). At *N* = 5 in Scenario 2, for example, mean total absolute estimation error for the Latino-like simulated admixed population decreased from 0.380 for the uncorrected estimates to 0.246 with the 1/*N* correction and 0.323 with the power-law *a*/*N*^*b*^ correction. Differences among the uncorrected and corrected estimates decreased as the reference sample size of the varying reference group increased.

### Practical guide for bias correction

Together, these results provide practical guidance for selecting a bias correction when reference sample sizes are small. When all reference groups have small reference sample sizes, as in Scenario 1, the power-law *a*/*N*^*b*^ correction generally provides the greatest reduction in total estimation error, particularly at small reference sample sizes. In contrast, when one reference group has a small reference sample size while the other reference groups have larger reference sample sizes, as in Scenarios 2 and 3, the 1/*N* correction provides more consistent reductions in total estimation error. Finally, once reference sample sizes are sufficiently large (above the practical threshold of N>150), both corrections provide minimal additional benefit, and uncorrected Summix2 is typically adequate.

**Table 1.** Practical guidance for bias correction in Summix2. Suggested corrections are based on correction performance across the reference sample-size scenarios evaluated in the simulation analyses. Performance was assessed using total absolute estimation error.

| Reference sample-size scenario | Suggested correction | When correction matters most | Rationale |
|---|---|---|---|
| **All reference groups have small reference sample sizes** | Power-law *a*/*N*<sup>*b*</sup>correction | When reference sample sizes are jointly limited across reference groups | Flexible; accounts for both the scale and decay of the relationship between estimation error and reference sample size |
| **One reference group has a small reference sample size while the others have larger reference sample sizes** | 1/*N* correction | When one reference group has a substantially smaller reference sample size than the others | Simple; does not require fitted parameters |
| **Reference groups have sufficiently large reference sample sizes** | Correction may provide limited additional benefit | Above the practical threshold (*N* > 150) | Avoids unnecessary correction |

### Real-data evaluation supports the simulation findings and identifies limitations of bias correction

To evaluate correction performance using real genetic summary data, we conducted stress tests across varying reference sample sizes. Because the true mixing proportions are unknown in real data, estimates obtained using the largest available reference sample sizes—*N* = 500 for AFR, EAS, EUR, and SAS and *N* = 45 for IAM—were used as a benchmark for stability rather than as ground truth.

In Setting 1, all five reference groups had the same reference sample size from *N* = 5 to *N* = 45. At larger evaluated reference sample sizes, IAM remained fixed at *N* = 45, while the reference sample sizes of AFR, EAS, EUR, and SAS increased to *N* = 500 (Figure 4A). Total absolute deviation from the benchmark was greatest at the smallest reference sample sizes. At *N* = 5, total absolute deviation decreased from 0.529 for the uncorrected estimates to 0.378 with the 1/*N* correction and 0.398 with the *a*/*N*^*b*^ correction. At *N* = 15, total absolute deviation decreased from 0.231 to 0.181 with the 1/*N* correction and 0.083 with the *a*/*N*^*b*^ correction. Both corrections also reduced total absolute deviation at *N* = 25, *N* = 35, and *N* = 45. However, at *N* = 75, *N* = 100, and *N* = 150, both corrections increased total absolute deviation relative to the uncorrected estimates. These results show that correction reduced total absolute deviation from the benchmark at the smallest reference sample sizes, whereas at several larger reference sample sizes, corrected estimates had greater total absolute deviation from the benchmark than uncorrected estimates.

**Figure 4A.**
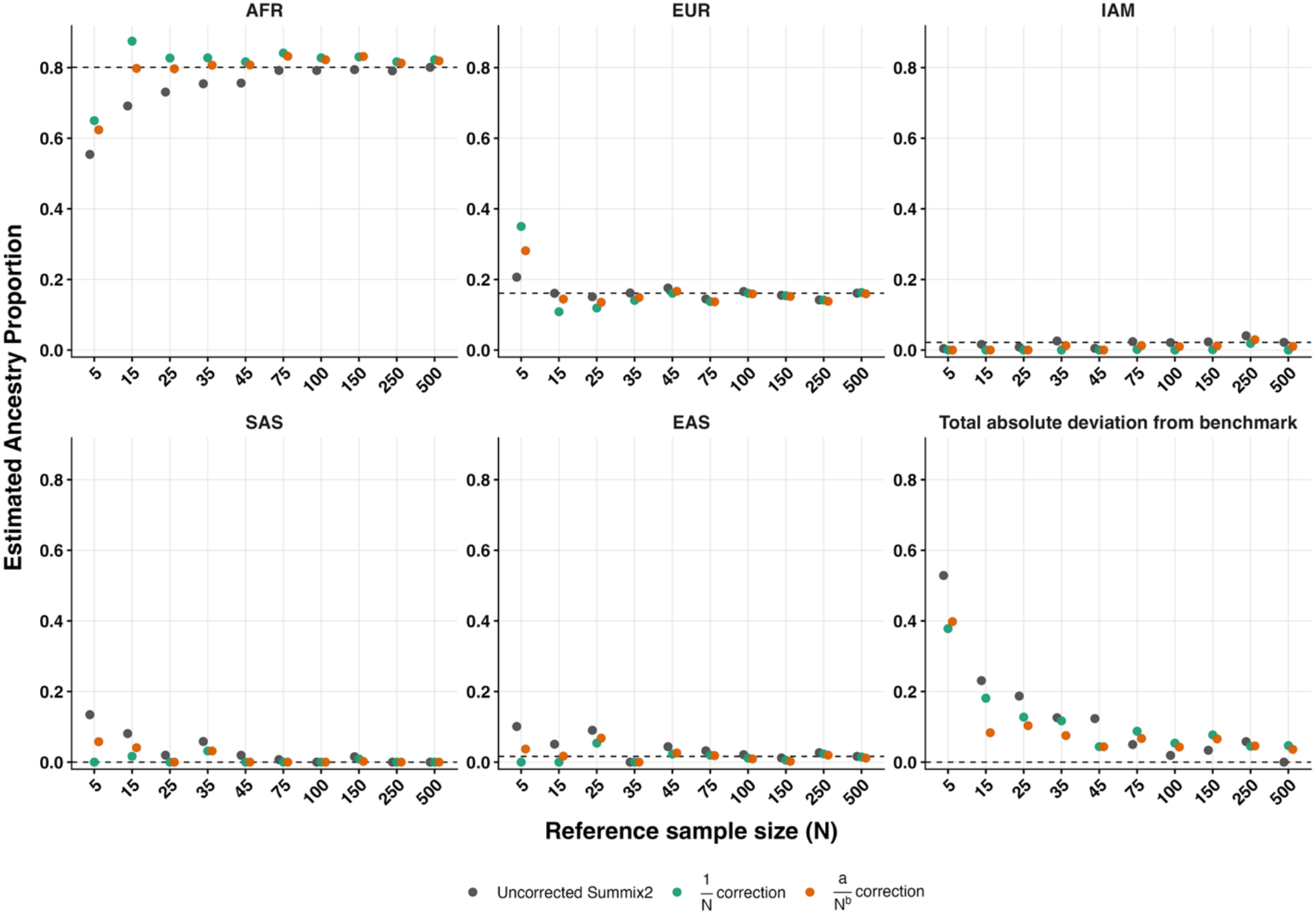
Real-data evaluation with reference sample sizes varied together where available. Mixing-proportion estimates are shown as a function of reference sample size for uncorrected Summix2 estimates and estimates after application of the 1/*N* and power-law *a*/*N*^*b*^ corrections. All five reference groups had the same reference sample size through *N* = 45. At larger evaluated reference sample sizes, IAM remained fixed at *N* = 45, while the reference sample sizes of AFR, EAS, EUR, and SAS increased. Dashed horizontal lines indicate benchmark estimates obtained using the largest evaluated reference sample sizes (*N* = 500 for AFR, EAS, EUR, and SAS and *N* = 45 for IAM). The bottom-right panel shows total absolute deviation from the benchmark across the five reference groups. The x-axis is shown on a log scale.

### Limiting the reference sample size of one reference group increases deviation across multiple reference groups in real data

In Setting 2, the AFR reference sample size was varied from *N* = 5 to *N* = 500, while the reference sample sizes of EAS, EUR, and SAS were fixed at *N* = 500 and IAM was fixed at *N* = 45 (Figure 4B). At an AFR reference sample size of *N* = 5, absolute deviation from the benchmark was 0.293 for AFR and 0.177 for EUR, despite the EUR reference sample size remaining fixed at *N* = 500. Total absolute deviation from the benchmark was 0.629 for the uncorrected estimates and decreased to 0.465 with the 1/*N* correction and 0.597 with the power-law *a*/*N*^*b*^ correction. As the AFR reference sample size increased, total absolute deviation generally decreased, reaching 0.0163 for the uncorrected estimates at *N* = 150, compared with 0.0143 for the 1/*N* correction and 0.0149 for the power-law *a*/*N*^*b*^ correction. Across the evaluated AFR reference sample sizes, the 1/*N* correction consistently produced the lowest total absolute deviation from the benchmark, although differences among methods became small as the AFR reference sample size increased. These results show that limiting the reference sample size of one reference group can increase deviation from the benchmark for other reference groups whose reference sample sizes remain fixed, consistent with the pattern observed in Scenarios 2 and 3.

**Figure 4B.**
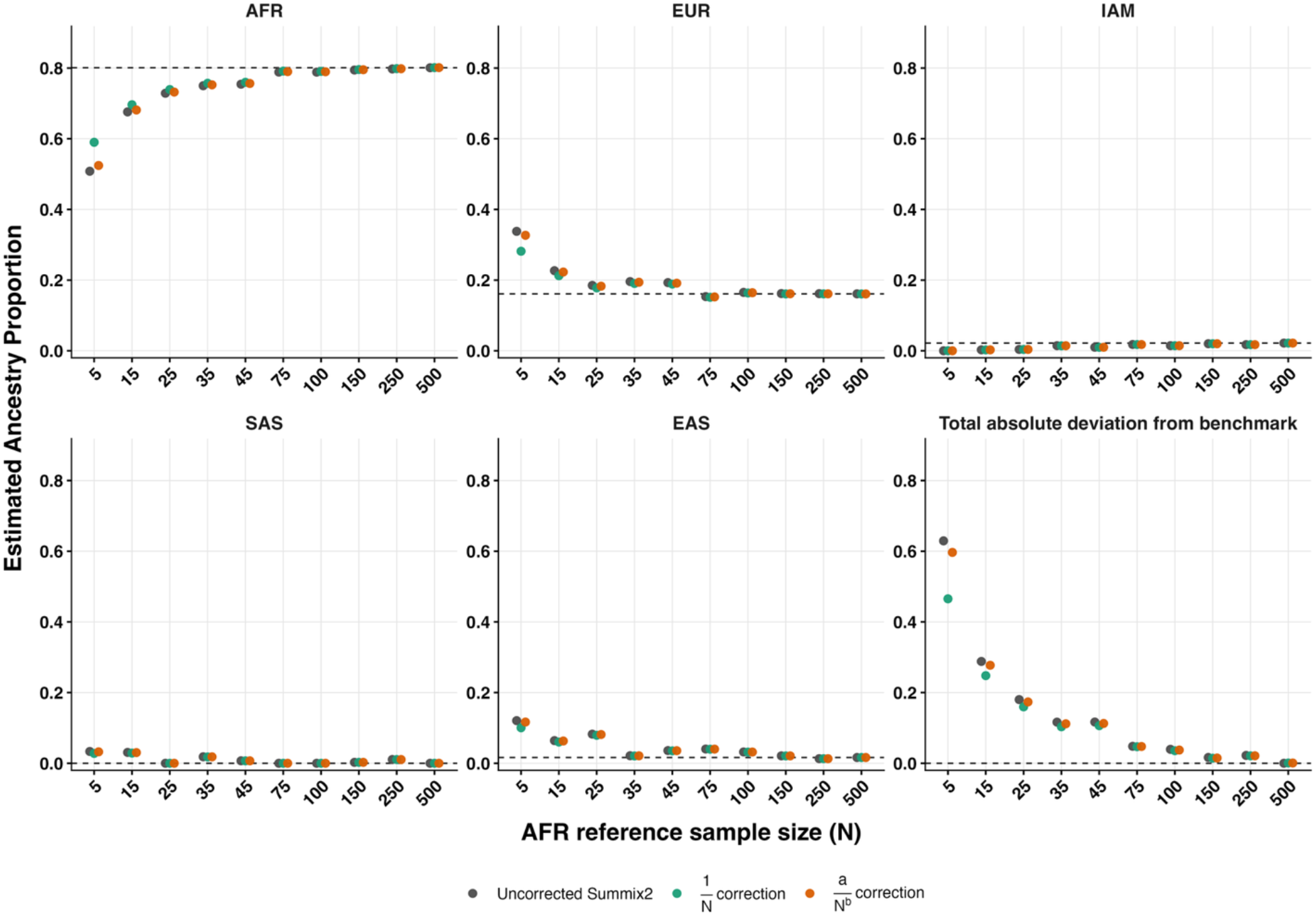
Real-data evaluation with the AFR reference sample size varied while the remaining reference sample sizes were fixed. Mixing-proportion estimates are shown as a function of AFR reference sample size for uncorrected Summix2 estimates and estimates after application of the 1/*N* and power-law *a*/*N*_*k*_ corrections. The AFR reference sample size was varied from *N* = 5 to *N* = 500, while the reference sample sizes of EAS, EUR, and SAS were fixed at *N* = 500 and IAM was fixed at *N* = 45. Dashed horizontal lines indicate benchmark estimates obtained using the largest available reference sample sizes (*N* = 500 for AFR, EAS, EUR, and SAS and *N* = 45 for IAM). The bottom-right panel shows total absolute deviation from the benchmark across the five reference groups. The x-axis is shown on a log scale.

## Discussion

We systematically evaluated the effect of reference sample size on Summix2 mixing-proportion estimation and tested two reference-sample-size corrections. Four main findings emerged. First, estimation error decreased as reference sample size increased, with the largest improvements occurring at small reference sample sizes. Second, across the simulation scenarios evaluated here, *N* = 150 emerged as a practical threshold, with relatively small reductions in estimation error at larger reference sample sizes. Third, limiting the sample size of one reference group increased estimation error in other reference groups whose sample sizes remained fixed. Finally, both the 1/*N* and power-law corrections reduced estimation error when reference samples were small, but their relative performance depended on how reference sample sizes were distributed across groups. Real-data analyses showed similar sensitivity to reference sample size but less benefit from correction when uncorrected estimates were already close to the large-reference benchmark.

Importantly, the effect of limited reference sample size was not confined to the reference group with limited data. In the Latino-like simulation, reducing the IAM reference sample size increased absolute estimation error for EAS even though the EAS reference sample size remained fixed at ***N*** = **500**. This pattern may reflect the joint estimation of constrained mixing proportions in Summix2, such that changes in the estimate for one reference group affect the estimates assigned to others. The pronounced effect on EAS may also reflect the similarity of the IAM and EAS reference allele frequencies, consistent with previous Summix findings, although we did not directly test this mechanism. These results demonstrate that the adequacy of a reference sample cannot be evaluated independently of the other reference groups included in the estimation.

More broadly, these findings show that reference sample size is an important consideration in population substructure estimation from genetic summary data. Reference allele frequencies are estimated from finite reference samples, and our results demonstrate that small reference sample sizes can increase estimation error not only for the reference group with limited sample size but also for other reference groups included in the estimation. Thus, the effect of reference sample size depends on the reference sample sizes across groups rather than on the sample size of each reference group independently. By characterizing these effects and evaluating approaches to reduce the resulting estimation error, this work provides practical guidance for population substructure estimation when reference sample sizes are small or unequal.

This study has several limitations. First, although our simulations were designed to be realistic, they necessarily simplify real population structure and do not capture all sources of heterogeneity present in empirical summary allele-frequency data, including finer-scale structure within continental reference groups. Second, the proposed approaches are post-estimation corrections and do not directly account for uncertainty in the reference allele frequencies used to estimate mixing proportions. Third, in the real-data analyses, performance was assessed relative to the large-reference benchmark rather than known mixing proportions. Because IAM reference data were limited, we were also unable to evaluate real-data settings in which all reference groups had similarly large sample sizes.

Several next steps follow from this work. One is methodological: the correction logic could be incorporated directly into the estimation procedure rather than applied after estimation. A second is robustness: future studies should evaluate how these corrections perform under reference mismatch and finer-scale substructure. A third is translational: it will be important to test how improved reference-size-aware correction affects downstream summary-data analyses, including applications such as PRS calibration and trans-ancestry meta-analysis correction. More broadly, the correction framework may extend to other reference-based estimation problems in genetics and genomics, particularly those that infer constrained mixture proportions from finite and potentially imbalanced reference data, such as cell-type deconvolution, local ancestry inference, and genotype or HLA imputation.

In summary, our findings demonstrate that reference sample size is an important consideration in Summix2 population substructure estimation and that the consequences of a small reference sample size can extend across reference groups. The effectiveness of post-estimation correction depends on how reference sample sizes are distributed across the reference groups included in the analysis, emphasizing the importance of considering the full reference composition rather than the sample size of an individual reference group in isolation. These findings provide practical guidance for applying Summix2 when reference data are limited and motivate future approaches that directly account for uncertainty in reference-group allele frequencies.

## Data and code availability

All data analyzed in this study were obtained from publicly available resources, including gnomAD, the 1000 Genomes Project, and the Human Genome Diversity Project, as described in the Methods. Code used to perform the simulations, apply the reference sample size-based corrections, conduct the real-data analyses, and generate the figures is available at https://github.com/Hendricks-Research-Team/summix2-bias-correction. The repository also includes the updated Summix2 implementation developed in this study.

## Acknowledgments

We gratefully acknowledge the publicly available data resources that made this study possible, including gnomAD, the 1000 Genomes Project, and the Human Genome Diversity Project. This work was supported by the National Human Genome Research Institute under award R35HG011293 and by the Pathway in Achieving Transferable Horizons in Genomic Data Science (PATH-GDS) program R25HG012919. This work also utilized computing resources at the Center for Computational Mathematics at the University of Colorado Denver, including the Alderaan cluster, supported by the National Science Foundation under award OAC-2019089.

## Author Contributions

S.T. and A.H. conceived the study. S.T. developed the methodology, implemented the simulation and bias correction workflows, performed the analyses, generated the visualizations, and wrote the manuscript. A.P. provided the data used for the simulation analysis and real-data evaluation. K.M. provided scientific mentorship and guidance on the population genetics aspects of the study. A.H. supervised the study and acquired funding. All authors reviewed and edited the manuscript and approved the final version.

## Declaration of generative AI and AI-assisted technologies in the writing process

During the preparation of this work, the authors used ChatGPT (OpenAI) to improve readability and language. After using this tool, the authors reviewed and edited the content as needed and take full responsibility for the content of the publication.

## References

1. Pasaniuc, B., and Price, A.L. (2017). Dissecting the genetics of complex traits using summary association statistics. Nature Reviews Genetics 18, 117–127. 10.1038/nrg.2016.142.

2. Sanderson, E., Glymour, M.M., Holmes, M.V., Kang, H., Morrison, J., Munafò, M.R., Palmer, T., Schooling, C.M., Wallace, C., Zhao, Q., and Smith, G.D. (2022). Mendelian randomization. Nat Rev Methods Primers 2. 10.1038/s43586-021-00092-5.

3. Kullo, I.J., Lewis, C.M., Inouye, M., Martin, A.R., Ripatti, S., and Chatterjee, N. (2022). Polygenic scores in biomedical research. Nature Reviews Genetics 23, 524–532. 10.1038/s41576-022-00470-z.

4. Wojcik, G.L., Murphy, J., Edelson, J.L., Gignoux, C.R., Ioannidis, A.G., Manning, A., Rivas, M.A., Buyske, S., and Hendricks, A.E. (2022). Opportunities and challenges for the use of common controls in sequencing studies. Nature Reviews Genetics 23, 665–679. 10.1038/s41576-022-00487-4.

5. Hendricks, A.E., Billups, S.C., Pike, H.N.C., Farooqi, I.S., Zeggini, E., Santorico, S.A., Barroso, I., and Dupuis, J. (2018). ProxECAT: Proxy External Controls Association Test. A new case-control gene region association test using allele frequencies from public controls. PLoS Genet 14, e1007591. 10.1371/journal.pgen.1007591.

6. Gouveia, M.H., Bentley, A.R., Leal, T.P., Tarazona-Santos, E., Bustamante, C.D., Adeyemo, A.A., Rotimi, C.N., and Shriner, D. (2023). Unappreciated subcontinental admixture in Europeans and European Americans and implications for genetic epidemiology studies. Nature Communications 14, 6802. 10.1038/s41467-023-42491-0.

7. Stoneman, H.R., Price, A.M., Trout, N.S., Lamont, R., Tifour, S., Pozdeyev, N., Anderson, H.D., Aquilante, C.L., Arbogast, K., and Arehart, C.H. (2025). Characterizing substructure via mixture modeling in large-scale genetic summary statistics. The American Journal of Human Genetics 112, 235–253.

8. Arriaga-MacKenzie, I.S., Matesi, G., Chen, S., Ronco, A., Marker, K.M., Hall, J.R., Scherenberg, R., Khajeh-Sharafabadi, M., Wu, Y., and Gignoux, C.R. (2021). Summix: a method for detecting and adjusting for population structure in genetic summary data. The American Journal of Human Genetics 108, 1270–1282.

9. Privé, F. (2022). Using the UK Biobank as a global reference of worldwide populations: application to measuring ancestry diversity from GWAS summary statistics. Bioinformatics 38, 3477–3480. 10.1093/bioinformatics/btac348.

10. Dennis, T., and Lee, D. (2024). ZMIX: estimating ancestry proportions using GWAS association Z-scores. Bioinformatics Advances 4. 10.1093/bioadv/vbae128.

11. Consortium, G.P. (2015). A global reference for human genetic variation. Nature 526, 68.

12. Carroll, S.R., Garba, I., Figueroa-Rodríguez, O.L., Holbrook, J., Lovett, R., Materechera, S., Parsons, M., Raseroka, K., Rodriguez-Lonebear, D., Rowe, R., et al. (2020). The CARE Principles for Indigenous Data Governance. Data Science Journal. 10.5334/dsj-2020-043.

13. Newman, M.E. (2005). Power laws, Pareto distributions and Zipf’s law. Contemporary physics 46, 323–351.

14. Chen, S., Francioli, L.C., Goodrich, J.K., Collins, R.L., Kanai, M., Wang, Q., Alföldi, J., Watts, N.A., Vittal, C., and Gauthier, L.D. (2022). A genome-wide mutational constraint map quantified from variation in 76,156 human genomes. BioRxiv, 2022.2003.2020.485034.

15. Cavalli-Sforza, L.L. (2005). The human genome diversity project: past, present and future. Nature Reviews Genetics 6, 333–340.

