## Supplementary Information for "A Bias Correction Method for Substructure Estimation with Small Reference Sample Sizes"

### Supplemental Information

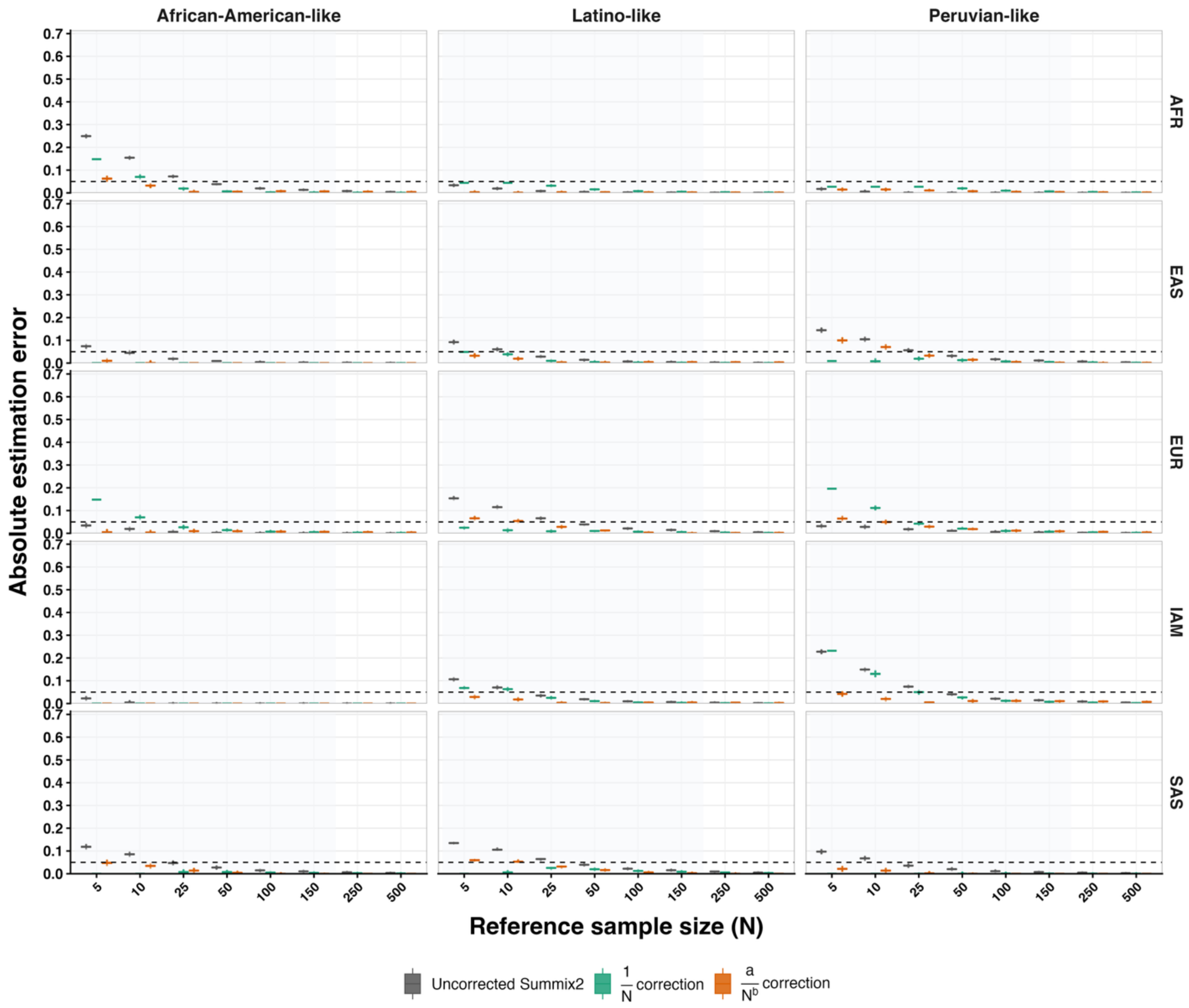


**Figure S1. Reference-group-specific absolute estimation error when all reference groups vary jointly with equal reference sample sizes (Scenario 1).** All five reference groups were assigned the same reference sample size at each evaluated value. Columns show the African American-like, Latino-like, and Peruvian-like simulated populations, and rows show the AFR, EAS, EUR, IAM, and SAS reference groups. Boxplots show the distribution of absolute estimation error, $|\hat{\pi}_k - \pi_k|$, across 1,000 simulation replicates for uncorrected Summix2, the $1/N$ correction, and the power-law correction. The dashed horizontal line indicates an absolute estimation error of 0.05, and shading denotes the low-reference regime ($N \leq 150$).

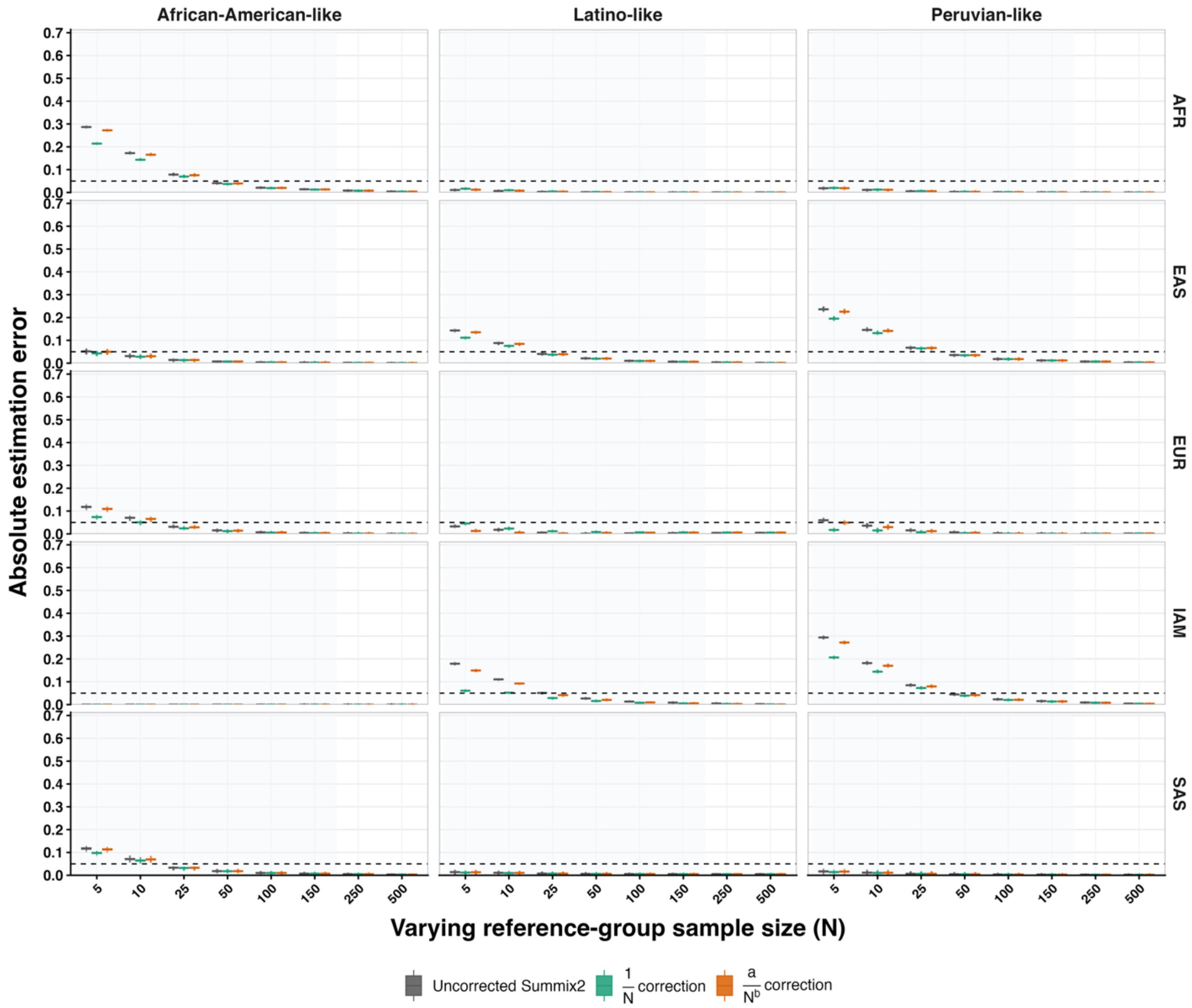


**Figure S2. Reference-group-specific absolute estimation error when one reference group varies and the remaining reference groups are fixed at $N = 500$ (Scenario 2).** The AFR reference sample size was varied for the African American-like population, whereas the IAM reference sample size was varied for the Latino-like and Peruvian-like populations; the other four reference groups were fixed at $N = 500$. Columns show the three simulated populations, and rows show the AFR, EAS, EUR, IAM, and SAS reference groups. Boxplots show the distribution of absolute estimation error, $|\ \hat{\pi}_k - \pi_k\ |$, across 1,000 simulation replicates for uncorrected Summix2, the $1/N$ correction, and the power-law correction. The dashed horizontal line indicates an absolute estimation error of 0.05, and shading denotes varying-reference-group sample sizes $N \leq 150$.

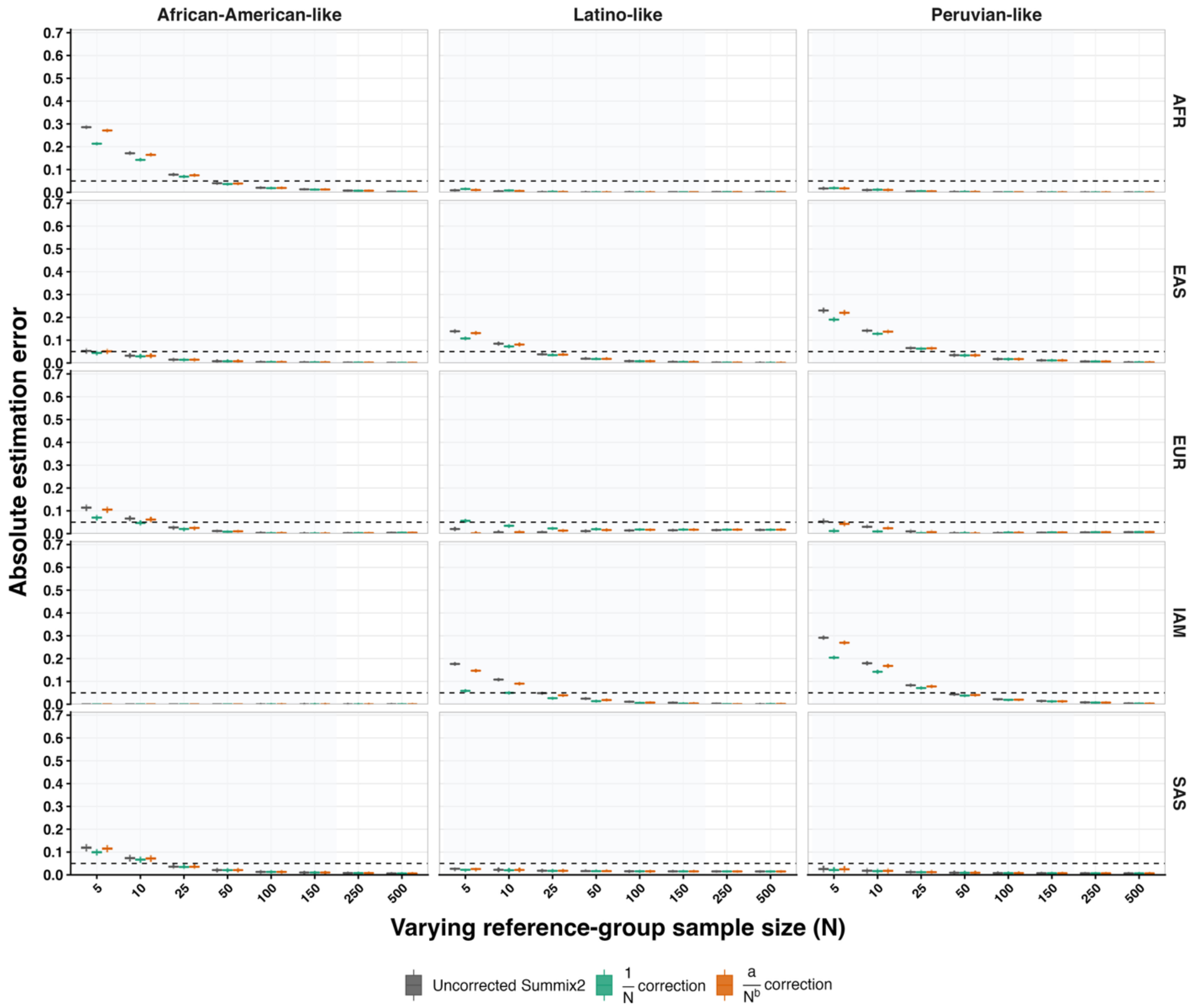


**Figure S3. Reference-group-specific absolute estimation error when one reference group varies and the remaining reference groups are fixed at $N = 150$ (Scenario 3).** The AFR reference sample size was varied for the African American-like population, whereas the IAM reference sample size was varied for the Latino-like and Peruvian-like populations; the other four reference groups were fixed at $N = 150$. Columns show the three simulated populations, and rows show the AFR, EAS, EUR, IAM, and SAS reference groups. Boxplots show the distribution of absolute estimation error, $| \hat{\pi}_k - \pi_k |$, across 1,000 simulation replicates for uncorrected Summix2, the $1/N$ correction, and the $a/N\hat{}b$ correction. The dashed horizontal line indicates an absolute estimation error of 0.05, and shading denotes varying-reference-group sample sizes $N \leq 150$.
